# First detection of High pathogenicity Avian Influenza A(H5N1) Clade 2.3.4.4b Genotype EA-2024-DI.2.1 in Egypt associated with migratory wild birds

**DOI:** 10.64898/2026.09.12.750890

**Authors:** Naglaa M. Hagag, Amany Adel, Mostafa R. Zaher, Ali Zanaty, Dalia Said, Asmaa Shaaban, Mostafa M. Saleh, Nehal M. Nabil, Maram M. Tawakol, Heba M. Hassan, Walaa I. Hussien, Naglaa Radwan, Chandana Tennakoon, Hamed Al-Aqnas, Samah Eid, Zienab Mosaad, Ahmed Samy

**Author notes:** Correspondence: Ahmed Samy (,) and Naglaa Haggag (]).

## Abstract

High pathogenicity avian influenza (HPAI) H5 clade 2.3.4.4b is the main driver of the ongoing unprecedented global panzootic. The recently emerged HPAI H5N1 clade 2.3.4.4b genotype EA-2024-DI.2.1 has become predominant in Europe, with migratory wild birds, particularly waterfowl, playing a major role in its dissemination. Egypt lies along major Afro-Eurasian migratory flyways, which have historically played an important role in the introduction of emerging H5Nx viruses into the country. In this study, targeted surveillance was conducted on 416 wild birds offered for sale in in live bird markets (LBMs) and roadside trading points in northern Egypt, mainly in Damietta and Port Said. Of these, 118 birds showing mild clinical signs were examined post-mortem and lung and tracheal tissues were collected, while oropharyngeal and cloacal swabs were collected from apparently healthy birds. Avian influenza virus was detected by RT-qPCR in 22 wild birds, all from tissue samples, whereas all swabs from apparently healthy birds were negative. Waterfowl accounted for 16 of the 22 positive birds (72.7%), with Eurasian teal showing the lowest Ct values (21-25). Phylogenetic and whole-genome analyses showed that the sequenced wild-bird viruses clustered within the recently emerged EA-2024-DI.2.1 sub-lineage and were closely related to contemporary European viruses. Compared with the EA-2021-AB genotype currently circulating in Egyptian poultry, the EA-2024-DI.2.1 viruses showed several HA amino acid differences, including A83D, L104M and T195A. These findings provide evidence for the introduction of EA-2024-DI.2.1 into Egypt through migratory wild birds and highlight the importance of continued genomic surveillance at the wild bird domestic poultry interface and antigenic evaluation against vaccines currently used in Egypt.

## Introduction

Highly pathogenic avian influenza (HPAI) H5N1 remains one of the most devastating livestock diseases globally due to its rapid evolution, global dissemination, massive impact on poultry production, and zoonotic potential. Influenza virus is an enveloped, negative-sense, single stranded RNA virus belongs to the *Orthomyxoviridae* family. The viral genome is formed of eight gene segments (PB2, PB1, PA, HA, NP, NA, M, and NS) each encoding 1-2 proteins with hemagglutinin (HA) and neuraminidase (NA) dictating antigenicity, as well as tropism, virulence and zoonotic potential (1–3). The HA protein has nineteen subtypes (H1–H19) and the NA protein has ten subtypes (N1–N10) that are mostly associated with wild birds (4).

The H5 A/Goose/Guangdong/1/1996 (gs/GD) lineage emerged in China in the 1990s and continued to evolute generating multiple HA clades, including clade 2.3.4.4 that emerged around 2014 that diversified into eight subclades (a–h), with clade 2.3.4.4b becoming the dominant lineage responsible for the ongoing global HPAI panzootic (5, 6). Continuous reassortment with low pathogenicity avian influenza viruses has driven remarkable genetic diversification with multiple genotypes disseminating globally mainly via wild birds. To date, more than 100 distinct reassortant genotypes within the H5Nx clade 2.3.4.4b have been identified worldwide. Importantly, many wild waterfowl species can carry clade 2.3.4.4b viruses with little or no clinical disease, facilitating viruses spread during migration (7, 8).

Egypt has experienced several major avian influenza crises since 2006 that have been linked to wild birds. This is related to Egypt’s location at the intersection of the Black Sea–Mediterranean and East Africa–West Asia migratory flyways, where millions of migratory waterfowl pass over the Nile Delta each year, making Egypt an important gateway for the movement of wild birds between Europe, Asia, and Africa (9). The HPAI H5N1 clade 2.2.1 virus was detected in a Eurasian teal at the beginning of the outbreak in December 2005 (10). Similarly, HPAI H5N8 was first detected in common coots and then in apparently healthy Eurasian teals in 2016 (11, 12). More recently, HPAI H5N1 clade 2.3.4.4b (EA-2021-AB genotype) was detected in a wild northern pintail and two domestic ducks in 2021 (6, 13). Notably, these incursions were first reported in northern Egypt, mainly in Port Said and Damietta, and were associated with wild birds offered for sale in LBMs, despite appearing clinically healthy.

In late 2024 in Europe, the HPAI H5N1 genotype EA-2024-DI became the predominant genotype, replacing the previous dominant genotype EA-2021-AB. EA-2024-DI contains five gene segments (PA, HA, NP, NA, and M) derived from the H5N1 EA-2021-AB genotype, while PB2, PB1, and NS originated from reassortment events, most likely with low pathogenicity avian influenza virus (LPAIV) circulating in wild Anseriformes in Eurasia. The DI genotype subsequently evolved into new sub-lineages, including EA-2024-DI.2.1, which was associated with a sharp increase in detections among wild waterfowl and with the majority of poultry outbreaks reported in Europe (14, 15).

To monitor potential new incursions of avian influenza viruses into Egypt, in the present study, we collected samples from migratory wild birds offered for sale in LBMs in northern Egypt. The sampling strategy included oropharyngeal swabs from apparently healthy birds and tissues from birds showing any apparent clinical signs. Samples were then subjected to molecular diagnosis and genetic characterization.

## 2. Materials and Methods

### 2.1. Sample Collection and Preparation

A total of 416 wild birds were sampled between September 2025 and April 2026 in northern coastal Egypt **(Supplementary Table S1)**. Wild birds were captured by local fishermen and, less frequently, local bird trappers using traditional trapping methods. The captured birds were subsequently offered for sale in LBMs or at roadside trading points along the main routes leading to Damietta, Port Said, and Ismailia. Oropharyngeal and cloacal swabs were collected from the sampled birds. Swabs from birds of the same species obtained from the same trader were pooled into a single sample, with a maximum of four birds per pool. Of the 416 sampled wild birds, 118 were purchased as whole birds to collected lung and trachea because they showed mild clinical signs as isolation from other birds, reduced activity or abnormal behaviour compared with other birds of the same species kept in the same cage. Both tissue homogenates and swab were thoroughly vortexed then centrifuged at 3,000 rpm for 15 min at 4°C, and the resulting supernatants were stored at −80°C until further analysis.

Samples collections and post-mortem examination procedures were reviewed and approved by the ethical committee at the Animal Health Research Institute, Ministry of Agriculture and Land Reclamation, Egypt.

### 2.2 Molecular diagnosis

All samples subjected to Viral RNA using the Patho Gene-spin™ DNA/RNA Extraction Kit (iNtRON Biotechnology, Seongnam, Republic of Korea) according to the manufacturer’s instructions. Nucleic acid extracts were screened by real-time RT-PCR (rRT-PCR) for the presence of the AIVs matrix (M) gene. Samples were also screened for Newcastle disease virus (NDV) and infectious bronchitis virus (IBV) to investigate potential co-circulation and mixed infections. Primers, probes, and thermal cycling conditions for all assays are provided in **Supplementary Table S2** (16–18). All RT-qPCR assays were carried out on an Applied Biosystems StepOnePlus™ Real-Time PCR System (Applied Biosystems, Foster City, CA, USA).

### 2.3 Virus Isolation and Hemagglutination Assay

AIV-positive samples identified by RT-qPCR were subjected to virus isolation using specific-pathogen-free (SPF) embryonated chicken eggs, following incubation, allantoic fluid was harvested and screened for hemagglutinating activity using hemagglutination (HA) assay using chicken red blood cells. Egg inoculation, harvesting and HA assay were performed according to the World Organization for Animal Health (WOAH) reference protocols (19).

### 2.4 Whole-genome sequencing and analysis

Whole-genome sequencing was performed using the primers listed in **Supplementary Table S3.** Viral gene segments were amplified by one-step RT-PCR using the EasyScript® One-Step RT-PCR Kit (TransGen Biotech, Beijing, China) in a ProFlex™ PCR System (Applied Biosystems, Waltham, MA, USA). The amplified products were separated by agarose gel electrophoresis and purified using the QIAquick Gel Extraction Kit (Qiagen, Hilden, Germany). Purified amplicons were subjected to Sanger sequencing using the BigDye™ Terminator v3.1 Cycle Sequencing Kit (Applied Biosystems). Sequencing reactions were purified using Centri-Sep™ spin columns and analysed on an ABI 3500xL Genetic Analyzer (Life Technologies, USA).

The retrieved sense and antisense sequences were assembled and trimmed, and a consensus was generated using UGENE and manually reviewed using BioEdit v7.2.5. Sequences generated in this study were deposited in the Global Initiative on Sharing All Influenza Data (GISAID) database under accession numbers EPI5609324–EPI5609343.

Representative global and Egyptian sequences representing different genotypes were retrieved from GISAID database. Sequence alignment was conducted using MEGA11 and BioEdit software. Neighbor-Joining phylogenetic trees were constructed using MEGA11 software, with parameters set to 1,000 bootstrap replications and the Maximum Composite Likelihood model. The trees were viewed using Fig tree V1.4.2 (http://tree.bio.ed.ac.uk/software/figtree/). Key amino acid mutations were identified using MEGA11 and BioEdit software and confirmed by relevant tools on GISAID. Potential N-linked glycosylation sites in the HA and NA proteins were predicted using the NetNGlyc 1.0 server (http://www.cbs.dtu. dk/services/NetNGlyc/) .

### 2.5. Genotype Determination of H5Nx Clade 2.3.4.4b Viruses

In addition to the phylogenetic analysis and confirmation using relevant tools on GISAID The genotype of H5Nx clade 2.3.4.4b viruses was confirmed based on the constellation of the eight viral gene segments (genotype backbone). Genotype assignment was performed using Genin2 software, which classifies viruses according to the genetic composition of their internal and surface gene segments (https://github.com/izsvenezie-virology/genin2?tab=readme-ov-file#input-guidelines).

## 3. Results

### 3.1. Molecular diagnosis

Using real-time RT-qPCR, avian influenza A virus was detected in 22 wild bird tissue samples, and all positive samples were negative for NDV and IBV. In contrast, none of the swabs collected from apparently healthy wild birds were positive for AIV.

Positive wild birds were detected between September and December 2025 in Damietta (Lake Manzala and Ras El-Bar) and Port Said (Table 1). The positive birds included Eurasian teal (n = 7), Northern Shoveler (n = 2), Northern pintail (n = 5), Peregrine falcon (n = 2), Greater flamingo (n = 2), Eurasian coot (n = 2), and Common pochard (n = 2). The lowest Ct values (21–25), indicating the highest viral RNA loads, were detected in Eurasian teal, while the other positive species generally showed higher Ct values (30–36) (Table 1).

**Table 1.**
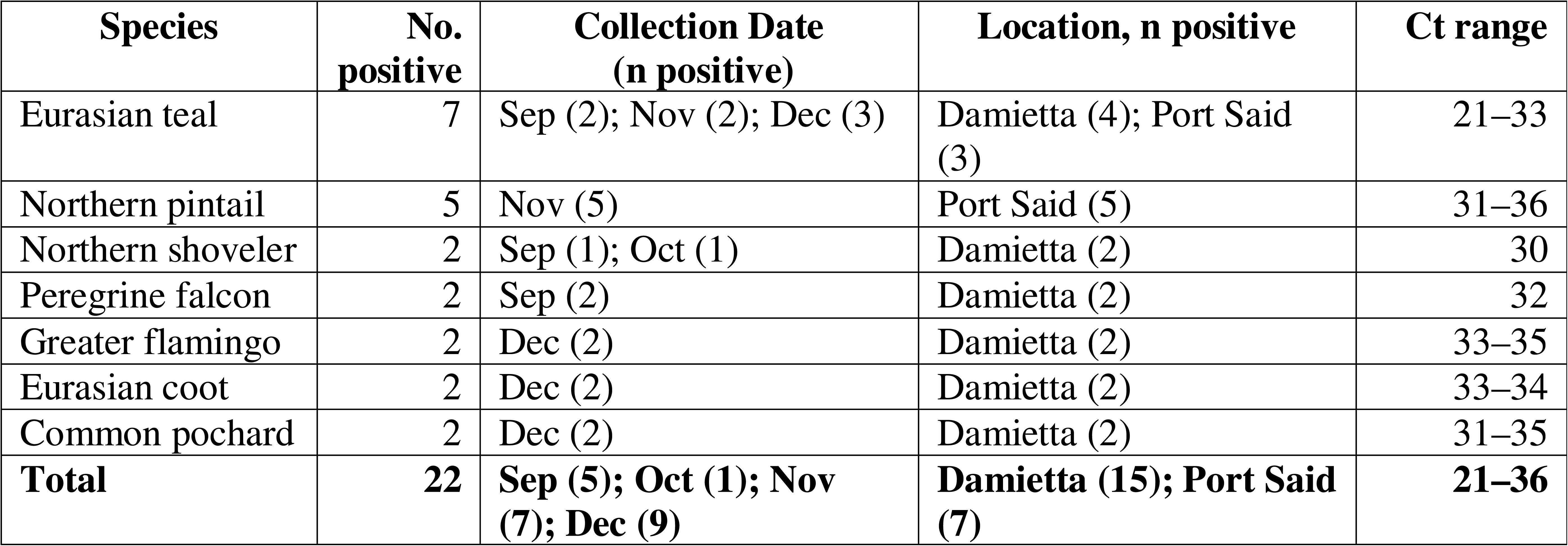
Species, temporal and geographical distribution of AIV-positive wild birds tissue samples:

### 3.2. Genetic characterization of the sequenced sample

#### HA gene

Phylogenetic analysis of the HA gene revealed clustering within the recently emerged EA-2024-DI.2.1 genotype detected in Germany, Bulgaria, Italy, and Spain (**Figure 1**) with around 99.4% nucleotide similarity. And around 97% nucleotide similarity with the endemic Egyptian EA-2021-AB genotype. Comparison of the deduced HA amino acid sequences identified several genotype-associated substitutions that distinguished the EA-2024-DI.2.1 viruses from the endemic EA-2021-AB genotype. Substitutions included A83D, L104M, T114I, P136S, K189N, and T195A. it worth mention that some previous Egyptian EA-2021-AB strains harbor T114I, P136S, K189N mutations. Among these, L104M is located within the receptor-binding site (RBS), whereas A83D, P136S, and T195A are located within antigenic sites E, A, and D, respectively. and T114I, and K189N are located within the globular head domain of HA (**Table 2**).

**Figure 1.**
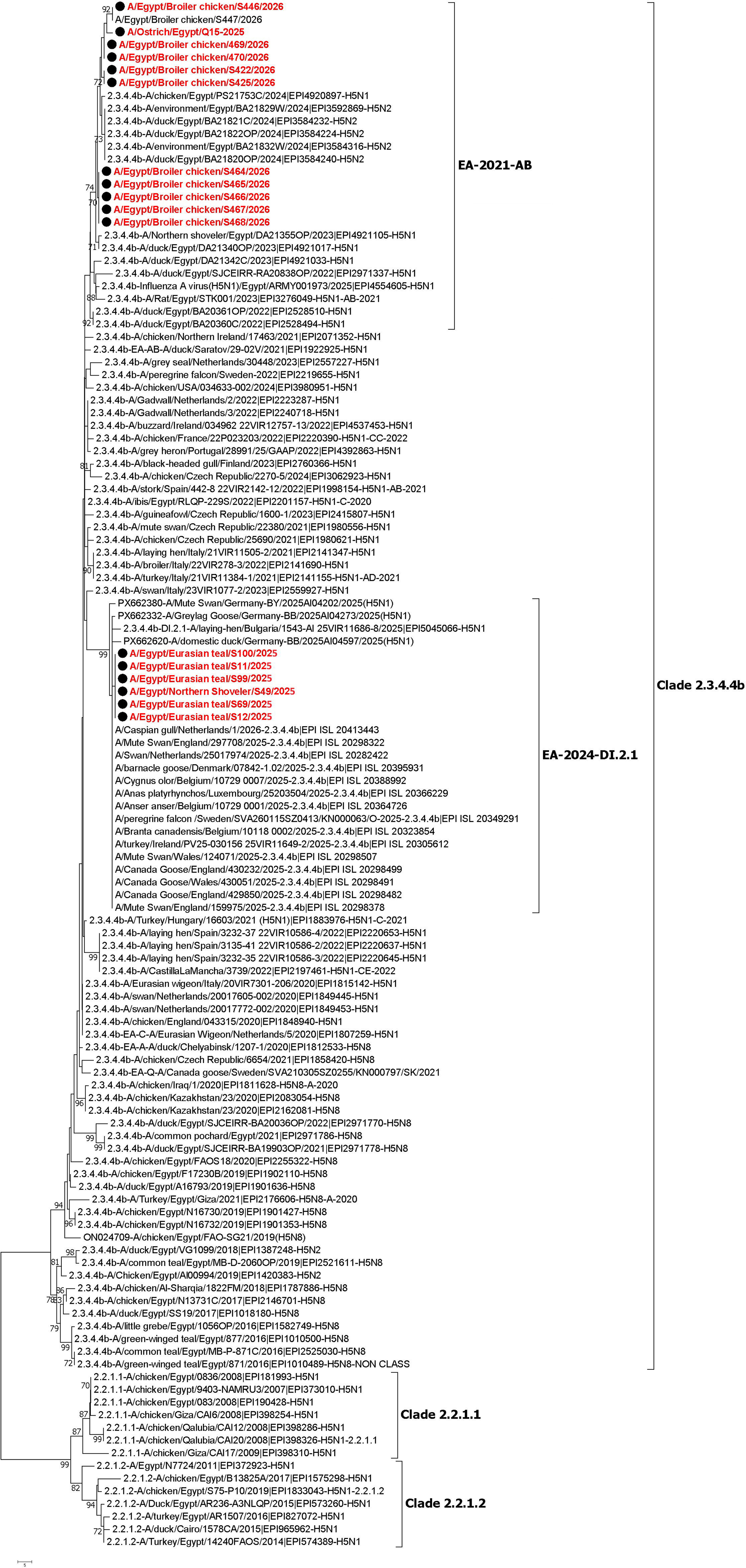
Phylogenetic analysis of the haemagglutinin (HA) gene of H5N1 clade 2.3.4.4b viruses. Viruses detected in wild birds in the present study are highlighted in red and clustered within the recently emerged EA-2024-DI.2.1 genotype, together with contemporary viruses detected in Germany, Bulgaria, Italy and Spain. Egyptian H5N1 sequences obtained from domestic poultry during the same period, but from different locations and independent of the wild-bird surveillance conducted in this study, were included to represent the H5N1 viruses currently circulating in Egyptian poultry. These viruses clustered within the EA-2021-AB genotype.

**Table 2:** the mutation analysis of HA amino acid sequence for the strains under investigation in comparison to the reference strains of the clade 2.3.4.4b.

| Strains | Genotype | RBS |  |  |  |  |  | mamalian markers |  |  | antigenic sites |  |  |  |  |  |  | Cleavage site |
| --- | --- | --- | --- | --- | --- | --- | --- | --- | --- | --- | --- | --- | --- | --- | --- | --- | --- | --- |
|  |  | 103 | 104 | 129 | 221 | 222 | 224 | 127 | 218 | 223 | 83 (E) | 114 (A) | 115 (A) | 136 (A) | 189 (B) | 195 (D) | 198 (D) | 323-332 |
| A/duck/Chelyabinsk/1207-1/2020_H5N8 | H5N8-EA-2020-A | H | L | L | G | Q | G | T | Q | R | A | I |  | P | N | T | I | REKRRKRGL<br>F |
| A/Canada_goose/Sweden/2021_H5N8 | H5N8-EA-2021-Q | H | L | L | G | Q | G | T | Q | R | A | I |  | P | N | T | I | REKRRKRGL<br>F |
| A/duck/Saratov/29-02V/2021_H5N1 | H5N1-EA-2021-AB | H | L | L | G | Q | G | T | Q | R | A | I |  | P | N | T | I | REKRRKRGL<br>F |
| A/Eurasian_Wigeon/Netherlands/5/2020_H5N1 | H5N1-EA-2021-C | H | L | L | G | Q | G | T | Q | R | A | I |  | P | N | T | I | REKRRKRGL<br>F |
| A/chicken/Egypt/PS21753C/2024_H5N1 | H5N1-EA-2021-AB | H | L | L | G | Q | G | T | Q | R | A | T |  | P | K | T | I | REKRRKRGL<br>F |
| A/Turkey/Egypt/Giza/2021_H5N8 | H5N8-EA-2020-A | H | L | L | G | Q | G | T | Q | R | A | I |  | P | N | T | I | REKRRKRGL<br>F |
| A/domestic duck/Germany/2025_H5N1 | H5N1-EA-2024-DI.2.1 | H | M | L | G | Q | G | T | Q | R | D | I | L | S | N | A | I | REKRRKRGL<br>F |
| A/EGY/Eurasian teal/S12/2025 | H5N1-EA-2024-DI.2.1 | H | M | L | G | Q | G | T | Q | R | D | I | L | S | N | A | I | REKRRKRGL<br>F |
| A/EGY/Eurasian teal/S99/2025 | H5N1-EA-2024-DI.2.1 | H | M | L | G | Q | G | T | Q | R | D | I | L | S | N | A | I | REKRRKRGL<br>F |
| A/EGY/Eurasian teal/S100/2025 | H5N1-EA-2024-DI.2.1 | H | M | L | G | Q | G | T | Q | R | D | I | L | P | N | A | I | REKRRKRGL<br>F |
| A/EGY/Eurasian teal/S11/2025 | H5N1-EA-2024-DI.2.1 | H | M | L | G | Q | G | T | Q | R | D | I | L | P | N | A | I | REKRRKRGL<br>F |

**Table 3:** The mutation analysis of NA amino acid sequence for the strains under investigation in comparison to the reference strains of the clade 2.3.4.4b

| Strains | Genotype | Deletion In<br>Stalk<br>Region | HB-sites |  |  | Oseltamivir<br>sensitive marker |  |  |  | N-glycosylation sites |  |  |  |  |  |  |  |  |  |
| --- | --- | --- | --- | --- | --- | --- | --- | --- | --- | --- | --- | --- | --- | --- | --- | --- | --- | --- | --- |
|  |  | (50-70 aa<br>residue) | 366-373 | 399-404 | 431-433 | 1<br>1<br>9 | 2<br>2<br>3 | 2<br>7<br>5 | 2<br>9<br>5 | 50 | 58 | 63 | 68 | 71 | 86 | 88 | 14<br>6 | 23<br>5 | 39<br>8 |
| A/duck/Chelyabinsk/1207-1/2020_H5N8 | H5N8-EA-2020-A | NO | TSRSG<br>FEI | WSG<br>YSG | PEE | E | I | H | N | - | NET<br>V | - | - | NT<br>SV | NN<br>TE | - | NG<br>TV | - | NW<br>SG |
| A/Canada_goose/Sweden/2021_H5N8 | H5N8-EA-2021-Q | NO | TSRSG<br>FEI | WSG<br>YSG | PEE | E | I | H | N | - | NET<br>V | - | - | NT<br>SV | NN<br>TE | - | NG<br>TV | - | NW<br>SG |
| A/duck/Saratov/29-02V/2021_H5N1 | H5N1-EA-2021-AB | NO | SSRSG<br>FEM | WSG<br>YSG | PKE | E | I | H | N | N<br>QS<br>I | NN<br>TW | NQ<br>TY | NI<br>SN | - | - | NS<br>SL | NG<br>TV | NG<br>SC | - |
| A/Eurasian_Wigeon/Netherlands/5/2020_H5N1 | H5N1-EA-2021-C | NO | SSRSG<br>FEM | WSG<br>YSG | PKE | E | I | H | N | N<br>QS<br>I | NN<br>TW | NQ<br>TY | NI<br>SN | - | - | NS<br>SL | NG<br>TV | NG<br>SC | - |
| A/laying-hen/Bulgaria/2025_H5N1 | H5N1-EA-2024-DI.2.1 | NO | SSRSG<br>FEM | WSG<br>YSG | PKE | E | I | H | N | N<br>QS<br>I | NN<br>TW | NQ<br>TY | NI<br>SN | - | - | NS<br>SL | NG<br>TV | NG<br>SC | - |
| PX662620-A/domestic duck/Germany-BB/2025_2025 | H5N1-EA-2024-DI.2.1 | NO | SSRSG<br>FEM | WSG<br>YSG | PKE | E | I | H | N | N<br>QS<br>I | NN<br>TW | NQ<br>TY | NI<br>SN | - | - | NS<br>SL | NG<br>TV | NG<br>SC | - |
| A/EGY/Broiler chicken/469/2026 | H5N1-EA-2021-AB | NO | SSRSG<br>FEM | WSG<br>YSG | PKE | E | I | H | N | N<br>QS<br>I | NN<br>TW | NQ<br>TY | NI<br>SN | - | - | NS<br>SL | NG<br>TV | NG<br>SC | - |
| A/EGY/Eurasian teal/S99/2025 | H5N1-EA-2024-DI.2.1 | NO | SSRSG<br>FEM | WSG<br>YSG | PKE | E | I | H | N | N<br>QS<br>I | - | NQ<br>TY | NI<br>SN | - | - | NS<br>SL | NG<br>TV | NG<br>SC | - |
| A/EGY/Eurasian teal/S12/2025 | H5N1-EA-2024-DI.2.1 | NO | SSRSG<br>FEM | WSG<br>YSG | PKE | E | I | H | N | N<br>QS<br>I | - | NQ<br>TY | NI<br>SN | - | - | NS<br>SL | NG<br>TV | NG<br>SC | - |

**Table 4:** the Pathogenic and mammalian markers for the strains under investigation in comparison to the reference strains of the clade 2.3.4.4b

| Strains | Genotype | M<br>1 | M2 |  | NP |  |  | NS |  |  |  |  | PA |  |  |  | PB<br>1 | PB1-F2 |  | PB2 |  |  |
| --- | --- | --- | --- | --- | --- | --- | --- | --- | --- | --- | --- | --- | --- | --- | --- | --- | --- | --- | --- | --- | --- | --- |
|  |  | 1<br>6<br>6 | 6<br>4 | 6<br>9 | 3<br>3 | 1<br>0<br>9 | 1<br>8<br>4 | 4<br>2 | 80–<br>84<br>Ami<br>no<br>Acid<br>Dele<br>tion | 9<br>2 | 1<br>4<br>9 | PD<br>Z<br>mo<br>tif<br>(22<br>7–<br>230<br>) | 1<br>2<br>7 | 5<br>5<br>0 | 6<br>1<br>3 | 672 | 13 | 7<br>3 | 82 | 6<br>2<br>7 | 5<br>0<br>4 | 7<br>0<br>1 |
|  |  | A | S | P | V | I | K | S | no | D | A<br>^ | GS<br>EV | V | L | E | L | P | K | S | E | V | D |
| A/Canada_goose/Sweden/2021 \| EPI<br>1853091-H5N8 | **H5N8-EA-**<br>**2021-Q** | - | - | - | - | - | - | - | no | - | - | GS<br>EV | - | - | - | - | - | - | - | - | - | - |
| A/duck/Saratov/29-02V/2021 \| EPI1922925-H5N1 | **H5N1-EA-**<br>**2021-AB** | - | - | - | - | - | - | - | no | - | - | <u>ES</u><br>EV | - | - | - | - | - | - | - | - | - | - |
| A/Eurasian_Wigeon/Netherlands/5<br>/2020 \| EPI1807259-H5N1 | **H5N1-EA-**<br>**2021-C** | - | - | - | - | - | - | - | no | - | - | <u>ES</u><br>EV | - | - | - | - | - | - | - | - | - | - |
| A/laying-hen/Bulgaria/2025 \| EPI5045066-H5N1 | **H5N1-EA-**<br>**2024-DI.2.1** | - | - | - | - | - | - | - | no | - | - | <u>EP</u><br>EV | - | - | - | - | - | - | - | - | - | - |
| PX662620-A/domestic<br>duck/Germany-BB/2025AI04597/2025 | H5N1-EA-2024-DI.2.1 | - | - | - | - | - | - | - | no | - | - | <u>ES</u><br>EV | - | - | - | - | - | - | - | - | - | - |
| **A/EGY/Eurasian teal/S99/2025** | **H5N1-EA-**<br>**2024-DI.2.1** | - | - | - | - | - | - | - | no | - | - | <u>ES</u><br>EV | - | - | - | - | - | - | P | - | - | - |
| **A/EGY/Eurasian teal/S12/2025** | **H5N1-EA-**<br>**2024-DI.2.1** | - | - | - | - | - | - | - | no | - | - | <u>ES</u><br>EV | - | - | - | - | - | - | P | - | - | - |

#### NA gene

Phylogenetic analysis showed that the two Eurasian teal isolates clustered with the recently emerged EA-2024-DI.2.1 genotype (**Figure 2**) and shared approximately 98.7% nucleotide identity with contemporary European H5N1 viruses detected in Germany during 2025, consistent with the genotype assignment obtained from the HA gene analysis. Comparison of the deduced NA amino acid sequences showed that none of the viruses contained deletions within the NA stalk region (amino acid residues 50–70), indicating that the stalk length remained conserved among both genotypes. However, the two EA-2024-DI.2.1 viruses possessed N58E and N59H substitutions, resulting in the loss of the predicted N-linked glycosylation motif at residues 58NNTW61, a feature absent from the endemic Egyptian EA-2021-AB viruses. While retained amino acid residues associated with susceptibility to neuraminidase inhibitors, including H275, R292, and N294 (N1 numbering), indicating the absence of established molecular markers associated with oseltamivir resistance. Analysis of additional functional residues demonstrated that the haemadsorption/second sialic acid binding site (HB/2SBS) region of the N1 proteins was conserved. Consistent with other contemporary H5N1 clade 2.3.4.4b viruses, the two isolates possessed the characteristic N1 substitutions T366S, I373M, and E432K relative to the N8 subtype. A complete comparison of NA amino acid substitutions between the viruses identified in this study, representative Egyptian H5N1 viruses, and selected reference strains is presented in **Supplementary Table S4**.

**Figure 2.**
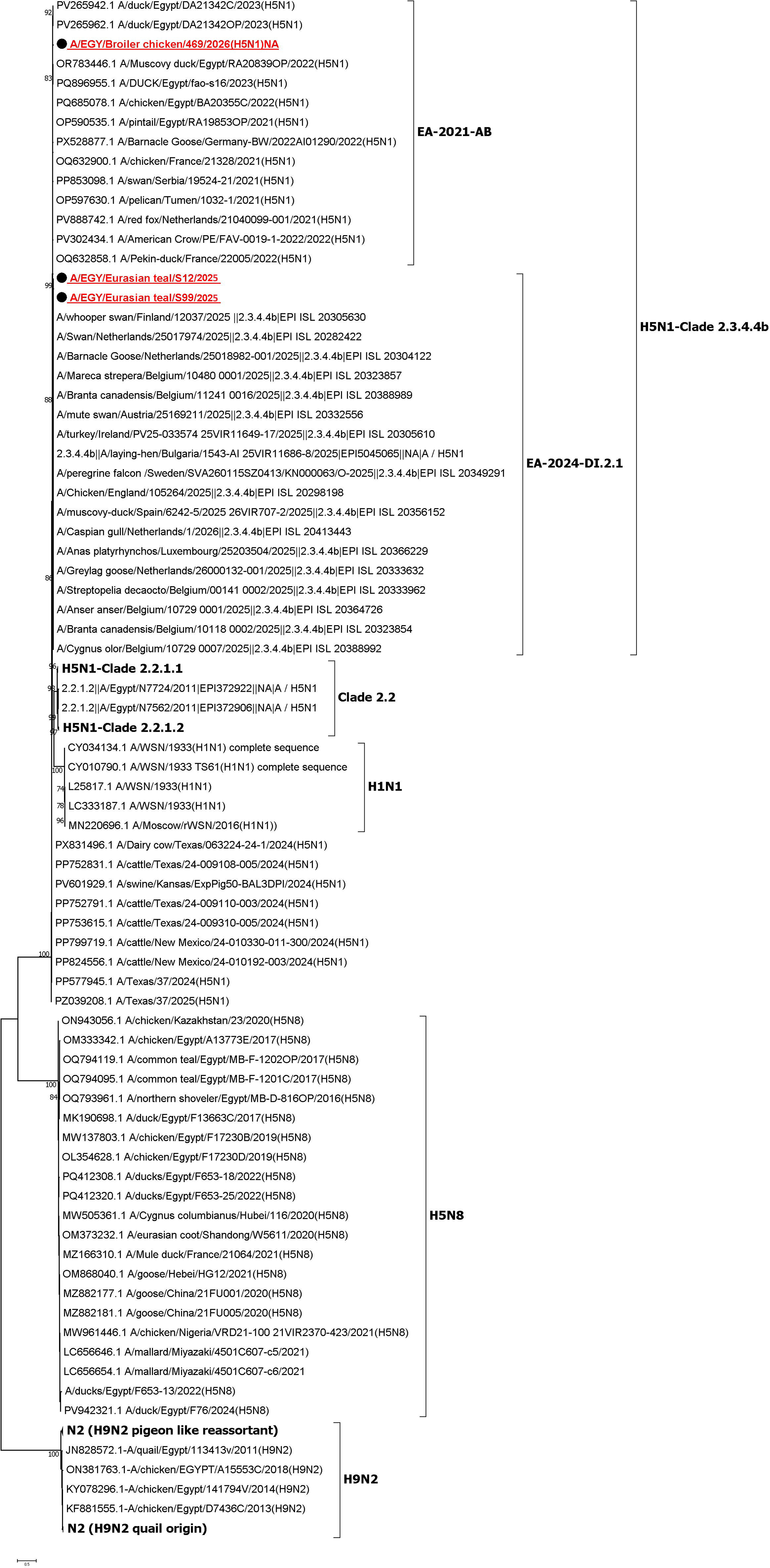
Phylogenetic analysis of the neuraminidase (NA) gene of H5N1 clade 2.3.4.4b viruses. Viruses detected in wild birds in the present study are highlighted in red and clustered within the recently emerged EA-2024-DI.2.1 genotype, together with contemporary viruses detected in Germany, Bulgaria, Italy and Spain. Egyptian H5N1 sequences obtained from domestic poultry during the same period, but from different locations and independent of the wild-bird surveillance conducted in this study, were included to represent the H5N1 viruses currently circulating in Egyptian poultry. These viruses clustered within the EA-2021-AB genotype.

#### Internal genes backbone

Genotyping analysis based on the complete internal gene sequences using the Genin2 classification tool confirmed that the two viruses recovered from migratory Eurasian teal (A/EGY/Eurasian teal/S12/2026 and A/EGY/Eurasian teal/S99/2026) were assigned to the recently emerged EA-2024-DI.2.1 genotype. These findings were consistent with the phylogenetic placement of the HA and NA gene analyses and suggesting a lack of further reassortment.

Phylogenetic analysis of the internal genes further supported these genotype assignments. The PB2, PB1, and NS genes of the two Eurasian teal viruses clustered with contemporary EA-2024-DI.2.1 viruses recently isolated from Europe, whereas their PA, NP, and MP genes clustered with EA-2021-AB lineage including the endemic EA-2021-AB lineage circulating in Egyptian poultry (**Figs. 3 and 4**).

**Figure 3.**
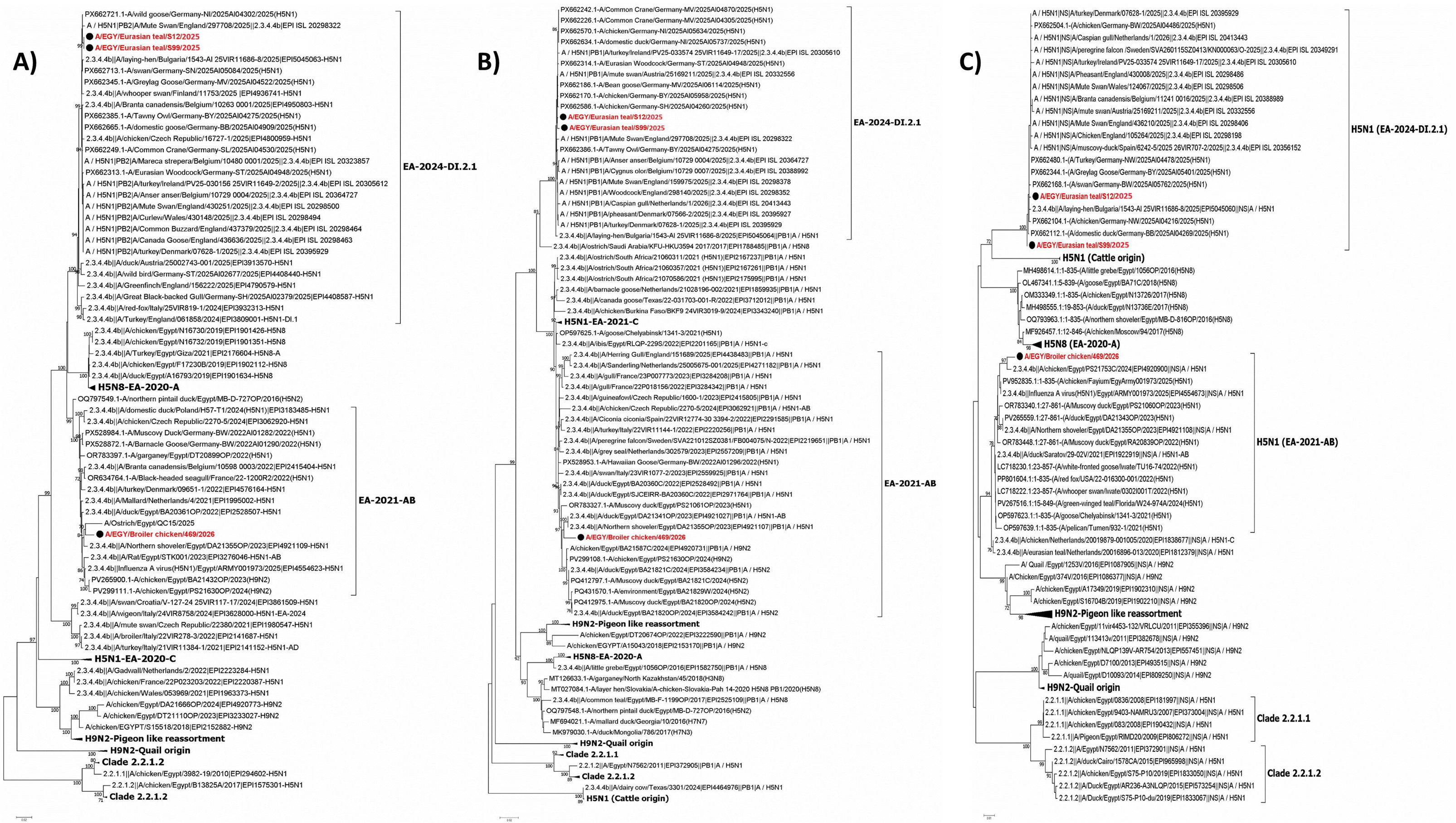
Phylogenetic analysis of the PB2, PB1 and NS genes of H5N1 clade 2.3.4.4b viruses. Viruses detected in wild birds in the present study are highlighted in red and clustered within the recently emerged EA-2024-DI.2.1 genotype, together with contemporary European viruses. Egyptian H5N1 sequences obtained from domestic poultry during the same period, but from different locations and independent of the wild-bird surveillance conducted in this study, were included to represent the H5N1 viruses currently circulating in Egyptian poultry. These viruses clustered within the EA-2021-AB genotype.

**Figure 4.**
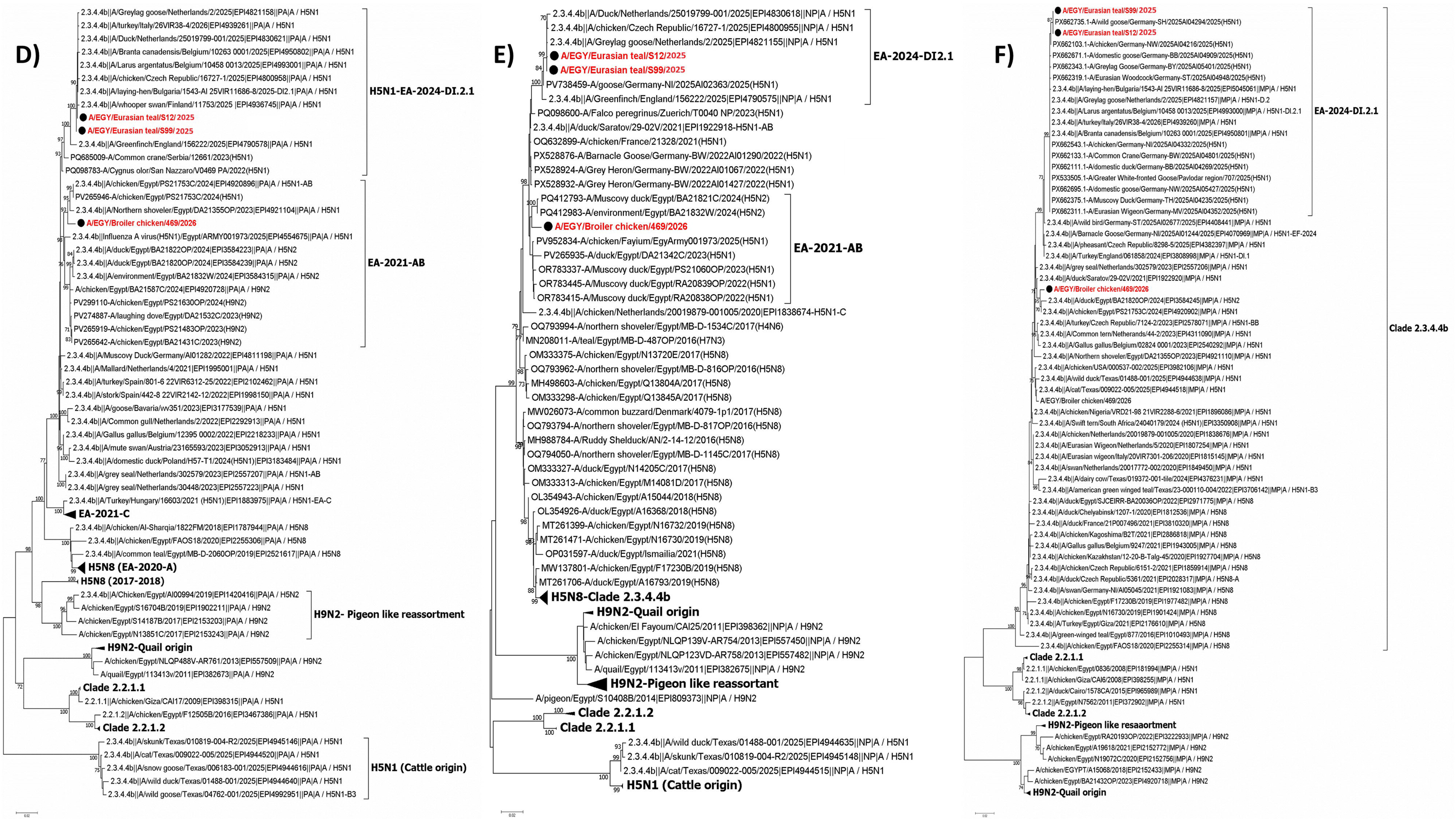
Phylogenetic analysis of the PA, NP and M genes of H5N1 clade 2.3.4.4b viruses. Viruses detected in wild birds in the present study are highlighted in red and clustered with contemporary EA-2024-DI.2.1 viruses. Egyptian H5N1 sequences obtained from domestic poultry during the same period, but from different locations and independent of the wild-bird surveillance conducted in this study, were included to represent the H5N1 viruses currently circulating in Egyptian poultry. In contrast to PB2, PB1 and NS, the PA, NP and M genes of EA-2024-DI.2.1 and EA-2021-AB share the same respective gene-segment lineages.

Comparison of the deduced amino acid sequences identified several molecular markers associated with viral pathogenicity, host adaptation, and antiviral susceptibility. The M2 protein retained the pathogenicity-associated residues S64 and P69 and lacked established markers associated with amantadine resistance, retaining valine at residue 27. Additional molecular signatures included S42 and an intact C-terminal PDZ-binding motif in NS1, together with V127, I550, and L672 in the PA protein. The PB2 protein retained glutamic acid at residue 627 (627E), consistent with an avian-adapted polymerase, while mammalian-associated markers P13 in PB1 and V504 in PB2 were also present. Furthermore, the PB1-F2 protein possessed the avian-associated residues K73 and S82P. A complete comparison of molecular markers identified in the viruses characterized in this study is presented in **Supplementary Table S5**.

## Discussion

The present study provides the first genomic evidence for the introduction of the recently emerged high pathogenicity avian influenza (HPAI) H5N1 clade 2.3.4.4b sub-genotype EA-2024-DI.2.1 into Egypt. To our knowledge, this represents the first report of EA-2024-DI.2.1 in Egypt and provides further evidence for the important role of migratory wild birds in the introduction of emerging H5Nx viruses into the country.

Wild aquatic birds are the natural reservoir of influenza A viruses and play a major role in the maintenance and long-distance dissemination of avian influenza viruses. In the present study, samples were collected from wild birds offered for sale LBMs in northern Egypt. All positive samples were detected in tissues collected from birds showing mild clinical abnormalities, whereas all swabs collected from apparently healthy wild birds were negative. Waterfowl accounted for 16 of the 22 positive wild birds (72.7%), including Eurasian teal, Northern shoveler, Northern pintail and Common pochard. Notably, Eurasian teal showed the lowest Ct values (21–25), indicating relatively high viral RNA loads. The predominance of infection among waterfowl is epidemiologically relevant, as these birds play an important role in the long-distance dissemination of avian influenza viruses during migration (9) . However, HPAI infection has also been reported in apparently healthy waterfowl, which may facilitate long-distance virus dissemination. Therefore, the mild clinical abnormalities observed in the positive wild birds in this study should be interpreted cautiously, as capture, transport and overcrowding under LBM conditions may also have influenced their clinical condition.

Phylogenetic analysis of positive samples revealed that all Egyptian wild bird viruses clustered within the recently emerged EA-2024-DI.2.1 sub-lineage circulating in Europe, where wild Anseriformes played a major role in virus spread (14). The EA-2024-DI genotype was first identified in wild Anseriformes in Eastern Europe in December 2023 and became the predominant genotype in Europe from October 2024. Unlike previous H5N1 epidemic waves, which were characterized by frequent reassortment events, the subsequent evolution of EA-2024-DI has so far been mainly driven by the gradual accumulation of mutations through genetic drift, leading to the emergence of the EA-2024-DI.1, EA-2024-DI.2 and, more recently, EA-2024-DI.2.1 sub-lineages. During the 2024–2025 epidemic, EA-2024-DI.2 became the predominant variant, accounting for 74% of genetically characterized viruses. At the beginning of the 2025–2026 epidemic, it was rapidly replaced across most of Europe by the EA-2024-DI.2.1 drift variant. Phylogenetic evidence suggests that EA-2024-DI.2.1 evolved from DI.2 viruses detected in Israel and the Republic of Georgia during the winter of 2024–2025, before its detection in Europe in September 2025 (14). Following its emergence, EA-2024-DI.2.1 spread rapidly across Europe and was associated with an unprecedented increase in HPAI detections in wild waterfowl during the 2025–2026 epidemic with numbers reaching six times those recorded during the previous two seasons (20) .

Interestingly, in the present study, EA-2024-DI.2.1 was detected in Egypt as early as September 2025, closely coinciding with its emergence in Europe. The Egyptian viruses clustered closely with European EA-2024-DI.2.1 viruses, suggesting a close epidemiological relationship. A recent study investigating the evolution of the H5N1 EA-2024-DI genotype in Europe estimated that the most recent common ancestor (tMRCA) of EA-2024-DI.2.1 dated to July 2025 (95% HPD: 6 June–11 August 2025). The sub-lineage was subsequently detected during the autumn 2025 migration, most likely introduced into Europe by migratory ducks arriving from breeding grounds in northern Eurasia (14). The detection of the same sub-lineage in migratory waterfowl in northern Egypt at approximately the same time raises the possibility that infected birds originating from common northern breeding areas contributed to introductions into both Europe and Egypt. However, introduction into Egypt following circulation through Europe cannot be excluded based on the available sequence data.

Like its ancestral EA-2024-DI genotype, EA-2024-DI.2.1 circulating in Europe and detected in the present study contains five gene segments (PA, HA, NP, NA and M) derived from the H5N1 EA-2021-AB genotype (5), while PB2, PB1 and NS originated from reassortment events, (14). In the present study, EA-2024-DI.2.1 viruses showed several HA amino acid differences compared with the EA-2021-AB genotype currently circulating in Egypt . However, the impact of these changes on antigenicity remains unclear. Sequence differences alone should not be interpreted as evidence of major antigenic drift or reduced vaccine protection. Further antigenic characterization, particularly cross-HI testing against vaccine strains currently used in Egypt, is therefore required to determine whether these substitutions affect antigenic matching and vaccine protection.

Importantly, the National Reference Laboratory for Poultry Diseases in Egypt conducts routine surveillance for avian influenza viruses in domestic poultry and regularly receives samples from different commercial poultry production systems across the country. At the time of writing, EA-2024-DI.2.1 has not been detected in domestic poultry through this surveillance. Therefore, there is currently no evidence of established circulation of this genotype in Egyptian domestic poultry. Several factors could contribute to this observation, including possible cross-protection provided by the widespread vaccination of poultry against H5N1 in Egypt. However, the current data are not sufficient to support this hypothesis. Continued genomic surveillance at the wild bird–domestic poultry interface, together with antigenic characterization and vaccine efficacy studies, will be important to determine whether currently used vaccines provide adequate protection against EA-2024-DI.2.1.

## Conclusion

In conclusion, the present study reports the detection of the recently emerged HPAI H5N1 EA-2024-DI.2.1 sub-lineage in Egypt, almost simultaneously with its detection and rapid spread in Europe. The virus was detected mainly in migratory waterfowl, and the Egyptian viruses formed a monophyletic cluster closely related to contemporary European EA-2024-DI.2.1 viruses, suggesting a potential common source associated with wild bird migration. Although EA-2024-DI.2.1 retains its HA gene from the EA-2021-AB genotype, several amino acid differences were identified compared with EA-2021-AB viruses currently circulating in Egypt. To date, EA-2024-DI.2.1 has not been detected in domestic poultry in Egypt. Continued molecular surveillance, particularly at the wild bird–domestic poultry interface, is therefore important to monitor potential spillover and further virus evolution. Antigenic characterization against vaccine strains currently used in Egypt is also required to determine whether existing vaccines provide adequate protection against this newly introduced sub-lineage.

## Acknowledgements

We would like to thank our local veterinary colleagues who facilitated sample collection, the fishermen who allowed us to sample their birds, and our colleagues at the Reference Laboratory for Veterinary Quality Control on Poultry Production for their technical assistance. We also thank Dr Thomas Peacock, The Pirbright Institute, for his valuable comments on the manuscript.

This work was funded by the British Council International Science Partnerships Fund (ISPF), UK, grant number 1203757062, and the Science, Technology & Innovation Funding Authority (STDF), Egypt, project ID 50185.

## References

1. Shaw M, Palese P. Orthomyxoviridae: The viruses and their replication. Fields virology. 2007;2.

2. Chen J, Lee KH, Steinhauer DA, Stevens DJ, Skehel JJ, Wiley DC. Structure of the hemagglutinin precursor cleavage site, a determinant of influenza pathogenicity and the origin of the labile conformation. Cell. 1998;95(3):409–17.

3. Gamblin SJ, Skehel JJ. Influenza hemagglutinin and neuraminidase membrane glycoproteins. J Biol Chem. 2010;285(37):28403–9.

4. Peiris JS, de Jong MD, Guan Y. Avian influenza virus (H5N1): a threat to human health. Clin Microbiol Rev. 2007;20(2):243–67.

5. Fusaro A, Zecchin B, Giussani E, Palumbo E, Agüero-García M, Bachofen C, et al. High pathogenic avian influenza A (H5) viruses of clade 2.3. 4.4 b in Europe—Why trends of virus evolution are more difficult to predict. Virus evolution. 2024;10(1):veae027.

6. Eid S, Hagag NM, Mosaad Z, Bakry NR, Elhusseiny MH, Mady WH, et al. Genomic surveillance and evolution of co-circulating avian influenza H5N1 and H5N8 viruses in Egypt, 2022-2024. Emerg Microbes Infect. 2025;14(1):2562046.

7. Couty M, Guinat C, Fornasiero D, Briand FX, Henry PY, Grasland B, et al. The role of wild birds in the global highly pathogenic avian influenza H5 panzootic, 2020-2023. NPJ Biodivers. 2026;5(1):1.

8. Chrzastek K, Lieber CM, Plemper RK. H5n1 clade 2.3. 4.4 B: evolution, global spread, and host range expansion. Pathogens. 2025;14(9):929.

9. Naguib MM, Verhagen JH, Samy A, Eriksson P, Fife M, Lundkvist Å, et al. Avian influenza viruses at the wild-domestic bird interface in Egypt. Infect Ecol Epidemiol. 2019;9(1):1575687.

10. Saad MD, Lu’ay SA, Gamal-Eldein MA, Fouda MK, Khalil FM, Yingst SL, et al. Possible avian influenza (H5N1) from migratory bird, Egypt. Emerging infectious diseases. 2007;13(7):1120.

11. Kandeil A, Kayed A, Moatasim Y, Webby RJ, McKenzie PP, Kayali G, et al. Genetic characterization of highly pathogenic avian influenza A H5N8 viruses isolated from wild birds in Egypt. Journal of General Virology. 2017;98(7):1573–86.

12. Selim AA, Erfan AM, Hagag N, Zanaty A, Samir A-H, Samy M, et al. Highly pathogenic avian influenza virus (H5N8) clade 2.3. 4.4 infection in migratory birds, Egypt. Emerging infectious diseases. 2017;23(6):1048.

13. El-Shesheny R, Moatasim Y, Mahmoud SH, Song Y, Taweel AE, Gomaa M, et al. Highly pathogenic avian influenza A (H5N1) virus clade 2.3. 4.4 b in wild birds and live bird markets, Egypt. Pathogens. 2022;12(1):36.

14. Zecchin B, Monne I, Dianati M, Bortolami A, Savegnago E, Shkodra E, et al. Two epidemics, one genotype, different outcomes: evolutionary changes of Avian Influenza H5N1, genotype EA-2024-DI. bioRxiv. 2026:2026.05. 25.727580.

15. Ducatez M, Fusaro A, Gonzales JL, Kuiken T, Ståhl K, Staubach C, et al. Unprecedented high level of highly pathogenic avian influenza in wild birds in Europe during the 2025 autumn migration. Efsa j. 2025;23(11):e9811.

16. Spackman E, Senne DA, Myers T, Bulaga LL, Garber LP, Perdue ML, et al. Development of a real-time reverse transcriptase PCR assay for type A influenza virus and the avian H5 and H7 hemagglutinin subtypes. Journal of clinical microbiology. 2002;40(9):3256–60.

17. Wise MG, Suarez DL, Seal BS, Pedersen JC, Senne DA, King DJ, et al. Development of a real-time reverse-transcription PCR for detection of Newcastle disease virus RNA in clinical samples. Journal of clinical microbiology. 2004;42(1):329–38.

18. Meir R, Maharat O, Farnushi Y, Simanov L. Development of a real-time TaqMan® RT-PCR assay for the detection of infectious bronchitis virus in chickens, and comparison of RT-PCR and virus isolation. Journal of virological methods. 2010;163(2):190–4.

19. Manual OT. Avian influenza (including infection with high pathogenicity avian influenza viruses. WOAH Terrestrial Manual. 2021:4.

20. Authority EFS, Prevention ECfD, Control, Influenza EURLfA, Barbezange C, Buczkowski H, et al. Avian influenza overview December 2025–February 2026. EFSA Journal. 2026;24(3):e10015.

